# ProteoParc: A tool to generate protein reference databases for ancient and non-model organisms

**DOI:** 10.1101/2025.07.31.667843

**Authors:** Guillermo Carrillo-Martin, Johanna Krueger, Tomas Marques-Bonet, Esther Lizano

## Abstract

Over the last few years, the increasing interest in analysing the proteome of extinct and non-model organisms has generated a new field of research expanding the scope of proteomics. The lack of curated databases and/or molecular data from these organisms forces researchers to manually search in different public repositories for related protein sequences, either for MS/MS peptide identification or ZooMS marker annotation. This can lead to format incongruences and hinder reproducibility between studies. To address this issue, we introduce ProteoParc, a user-friendly software that generates reference databases by systematically downloading and processing protein sequences from the most widely used public repositories. The pipeline’s output is a non-redundant protein database, formatted to be interpreted by typical peptide identification software. Moreover, the user can adjust the database dimension and composition by applying different criteria to include only a certain number of genes or species. Thus, ProteoParc is an easy and fast, custom-made bioinformatic tool useful for future paleoproteomics analysis in ancient samples related to understudied organisms.

## Introduction

It is well known that biomolecules can remain in the ecosphere for up to millions of years **(Cappellini et al., 2018)**. Although palaeogenetics has been proven to be a productive discipline for studying the past **(Dalén et al., 2023)**, DNA is unstable and rapidly degraded or banished in post-mortem conditions by temperature and microbial communities **(Brunson & Reich, 2019)**. By contrast, proteins are generally smaller and more compact, reducing their susceptibility to chemical degradation **(Warinner et al., 2022)**. While the oldest sequenced aDNA belongs to exceptionally well-preserved sediment samples dated approximately 2 million years (Ma) **(Kjær et al., 2022)**, recent studies have identified ancient protein sequences dating back to 21–24 Ma **(Paterson et al., 2025)**. In this sense, paleoproteomics has emerged as a molecular science discipline that aims to recover, identify, and analyse peptides in ancient tissues or biological remains **(Hendy et al., 2018)**. In the last decade, protein analysis through shotgun tandem mass spectrometry (MS/MS) helped to unravel some aspects of long-established evolutionary areas, such as palaeontology or archaeology, e.g. to solve the phylogeny of extinct species **(Cappellini et al., 2019; Presslee et al., 2019; Welker et al., 2019, 2020)**, perform chromosomal sexing on fossil individuals **(Mikšík et al., 2023)**, or study ancient human behaviour, from Palaeolithic **(M. Ma et al., 2024)** to recent historical events **(Di Gianvincenzo et al., 2023; Mackie et al., 2018)**. Additionally, the identification of species through zooarchaeology by mass spectrometry (ZooMS), a collagen fingerprinting technique, has been established as a powerful workflow for taxonomic assessment in bone remains **(Richter et al., 2022)**. The growth of genomics and the price reduction of genome sequencing have also enabled the possibility of analysing the proteome of non-model organisms, that is, the vast majority of species that have not been systematically studied. The applications of this horizon range from biomedical advances to new biological discoveries **(Heck & Neely, 2020)**.

Reference protein databases are a key aspect of mass-spectrometry-aided peptide analyses, as they serve as the source for: 1) in ZooMS, identifying the sequence behind a specific marker for its annotation in the standard nomenclature **(Brown et al., 2021)**; and 2) in shotgun MS/MS proteomics, generating theoretical mass-to-charge spectra to match experimental data and infer amino acid sequences **(Steen & Mann, 2004)**. In MS/MS tissue-specific analyses, the size and composition of these databases heavily affect the identification output. Large-database approaches, covering sequences of many species or tissues, raise the computational cost and likelihood of identifying false positives when using peptide identification software **(Rodriguez Palomo et al., 2024)** –e.g. MASCOT **(Perkins et al., 1999)**, SEQUEST **(MacCoss et al., 2002)**, PEAKS **(B. Ma et al., 2003)**, pFind **(Li et al., 2005)**, MaxQuant **(Cox & Mann, 2008)**, Byonic **(Bern et al., 2012)**, MsFragger **(Kong et al., 2017)**, MetaMorpheus **(Solntsev et al., 2018)**, or Proteome Discoverer **(Orsburn, 2021)**–. Thus, narrow databases identify peptides with higher statistical confidence, but sequences not present in the database will most likely not be detected, resulting in a set of unidentified experimental spectra **(Welker, 2018)**. In shotgun palaeoproteomic analyses, this is the norm more than the exception, as ancient peptides are potentially different to the sequences present in reference databases due to unpredictable evolutionary changes. Particularly, a recent meta-analysis revealed that more than 94% of the experimental spectra remain unidentified in this context (Chiang, Welker, et al., 2024).

Nowadays, narrow databases for ancient and non-model proteomics contain proteins known to be expressed in the tissue sample type. In the specific case of paleoproteomics, sequences are taken from an extant sister clade of the sampled extinct species **(Taurozzi et al., 2024)**. As many repositories must be consulted (e.g. UniProt **(The UniProt Consortium, 2025)** or RefSeq **(O’Leary et al., 2016)**), the manual process of building a database is repetitive and time-consuming, leading to format incongruences and missing data; these issues complicate any downstream analysis and may lead to errors in interpretation. Moreover, this process must be repeated each time a new species and tissue combination is analysed or even within the same study to optimise the reference database composition.

The increased interest in ancient and non-model protein studies makes it likely that more taxa will be studied under these methodologies. Thus, a solution to automatically generate custom and reproducible reference protein multi-FASTA is essential for efficiency and consistency. To address this issue, we propose ProteoParc, an easy-to-use software tool to generate optimised reference databases by systematically downloading and processing protein sequences from the most widely used public repositories.

## Methods

### Software Overview

ProteoParc is coded as a bioinformatic pipeline, concatenating scripts written in Python and R code languages (github.com/guillecarrillo/proteoparc). Its execution relies on a Python script (proteoparc.py), which any Linux-like terminal can execute. In short, the protein sequences download mechanism is based on iteratively searching records at UniParc (uniprot.org/help/uniparc), a non-redundant archive that stores peptide sequences from more than 20 repositories. The search scope is focused on a particular clade, specified by its NCBI TaxID. This ID is unique to each described taxon and shared between public sequence repositories. Alternatively, the search can be more specific if a gene list is input, which results in downloading only the proteins annotated under the genes in the list. This way, the database specificity increases, reducing downstream computational times and increasing the statistical confidence of peptide identifications in MS/MS analysis. After the database is built, records might be processed under two optional operations: a) remove redundant sequences to reduce extra comparisons by MS/MS identification software, or b) generate an aligned version of the recovered proteins to easily detect incomplete and low-quality sequences in manual inspection by the user. To summarise the metadata information, ProteoParc parses the final database version; results are output in a collection of tables, showing the number of genes, species, and repositories represented. Furthermore, a set of plots is generated to visually inspect for missing proteins or species not present in the repositories.

Although its use can be diverse, ProteoParc is intended to generate reference databases for non-model organisms and ancient protein analysis. As previously discussed in this context, it is crucial to include protein information from various extant and extinct species. This relates to either generating input databases for MS/MS identification software or building collagen alignments to analyse ZooMS markers among different species.

### Workflow Design

ProteoParc’s execution is divided into three main steps: Download, Processing and Metadata **(Figure 1A)**; Each of them is consecutively executed once the previous process has finished. They can also be run separately by executing the scripts in individual runs, allowing manual curation between steps. Two inputs, Project name and TaxID, are mandatory, while a list of genes is optional. Without a gene list, the whole clade proteome is downloaded. The output location may be modified, with the working directory being the default output path. The user can also deactivate the *Remove Redundancy* or *Alignment* processes before the execution to reduce computational time (see *Execution and Time Performance*).

**Figure 1.**
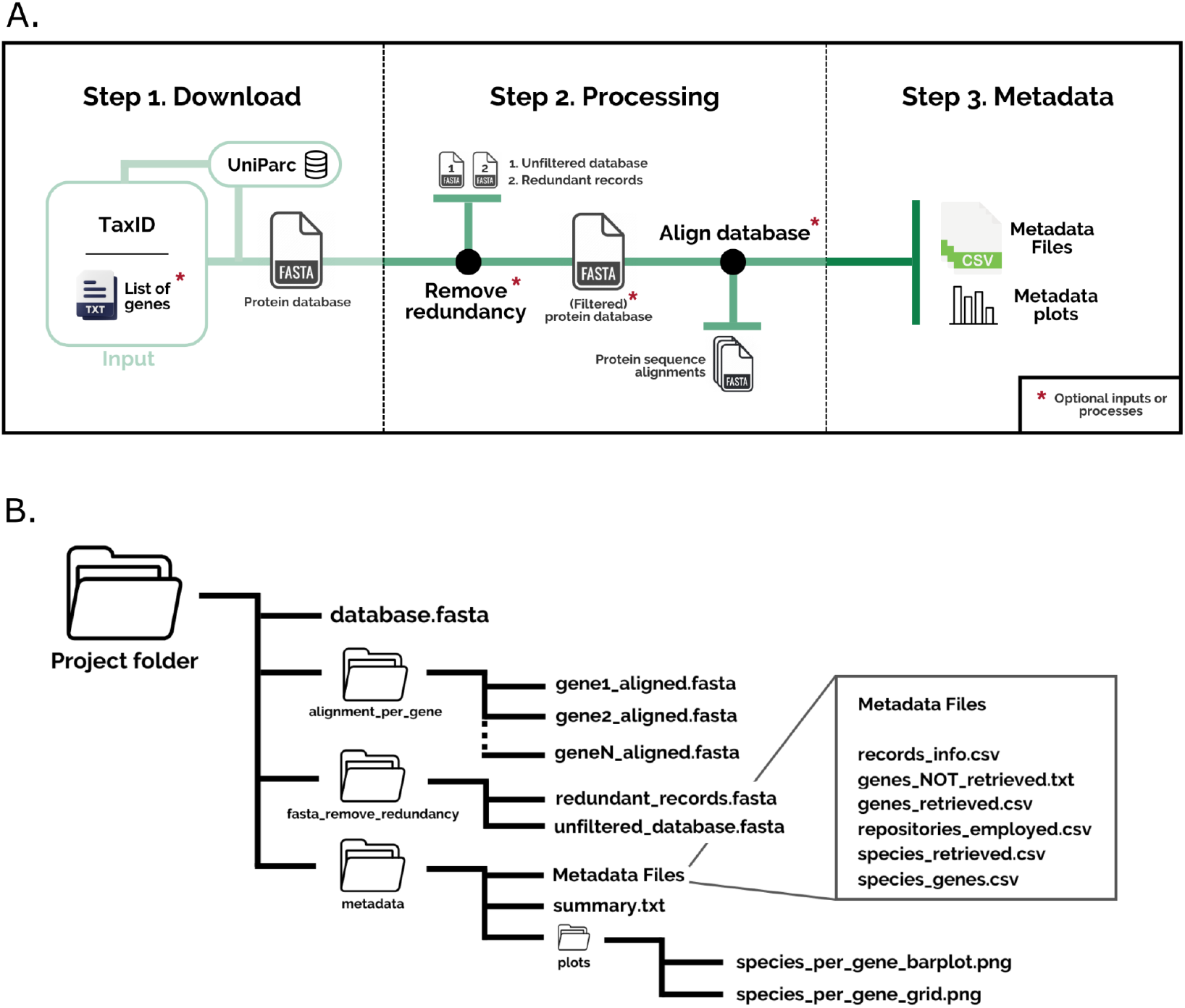
**A)** ProteoParc’s workflow overview. (1) Protein records are downloaded from UniParc under a specific TaxID and an optional list of genes, then written into a multi-FASTA file. (2) Duplicated and fragmentary sequences can be filtered out, and an aligned version of each protein in the multi-FASTA can be generated for manual curation purposes. (3) Finally, metadata information is collected from the multi-FASTA database and presented as a set of tables and plots. Optional inputs or processes are marked with an asterisk. **B)** Output display of a run with all the processes being activated: (1) the multi-FASTA protein database (database.fasta); (2.1) a folder containing the removed redundant records and the unfiltered database (fasta_remove_redundancy); (2.2) a folder containing the protein sequence alignments (alignment_per_gene); (3) a folder containing the metadata information as a set of tables and plots (metadata).

#### 1. Download step

A multi-FASTA database is built from all protein isoforms and variants that fulfil each search requirement, i.e the combination of a gene name and/or a TaxID. Proteins are downloaded through an informatics procedure, i.e application programming interface (API), which extracts information from a software component, in this case, UniParc **(Ahmad et al., 2025)**. The query search parameters are detailed within a URL link; If a gene list is input, the search process is repeated, including a different gene name within the URL query for each iteration. As a sequence can be associated with more than one species and repositories, three JSON files are generated, annotating the species, taxID and source repositories of each record. After the download, all headers are rearranged into the same format, including information about the source repository, UniParc’s ID, last update, protein name, species name, species TaxID, gene name and sequence version.

#### 2. Processing step

Two optional processes might be performed using the multi-FASTA database as input:

*a) Remove redundancy*: Although UniParc is a non-redundant archive, proteins can be downloaded twice if they are annotated under two different gene names present in the gene list. Thus, redundant proteins present in two or more different records are removed until only one copy remains. This process also removes sequences that are fragments (substrings) of other records in the database; For instance, short protein isoforms or incomplete sequences.

*b) Alignment*: An aligned version of the database is generated for each annotated gene. Proteins are first sorted based on their gene name and then aligned using mafft v7.525 **(Katoh & Standley, 2013)** under the --auto argument, which automatically selects the optimal strategy according to the sequence size. Records without a gene name will not be aligned through this process.

#### 3. Metadata step

Metadata information is extracted from the multi-FASTA record headers and presented as a set of tables, text files, and plots. This information is completed with the JSON dictionaries for those records that are associated with more than one species or source repository. However, this option can be disabled to just rely on the metadata within the multi-FASTA. In any case, JSON files are removed after the metadata has been extracted. The metadata tables summarise the number of retrieved genes, species and repositories, in addition to the number of genes retrieved per species. On the other hand, plots are generated to visualise the content of the database easily. All information related to the database name, download date, and metadata is condensed into a summary text file. Warning messages are printed there when records have missing information or if genes present in the gene list are not found.

## Results

### Output & Formatting

ProteoParc’s output is structured in three categories regarding the steps underlying the pipeline **(Figure 1B)**:

1) multi-FASTA protein database - *Download step*; All the records are parsed and formatted similarly, based on the UniProt structure. The metadata present in each header consists of information about the source origin, UniParc ID, last update, protein name, species, TaxID, and gene name. The protein name and gene name might be omitted in cases when a gene list is not input, as some proteins in the whole species proteome might not be properly annotated. Commas (,), semicolons (;) or colons (:) are removed, if present, to avoid interference with CSV formatting.

2.1) Redundancy FASTA files (optional) - *Processing step*; A filtered multi-FASTA database (without redundant sequences) substitutes the prime database version. Both the unfiltered database (unfiltered_database.fasta) and a file storing the removed records (redundant_records.fasta) are output in the fasta_remove_redundancy directory.

2.2) Protein sequence alignments (optional) - *Processing step*; All proteins in the database annotated under the same gene will be aligned and output as a multi-FASTA file with hyphens (“-”) representing gaps. All the alignments are stored in the alignment_per_gene directory, under the name x_aligned.fasta (*x* being a specific gene name).

3) Metadata information - *Metadata process*; six metadata files are output displaying:

I) count of retrieved genes (genes_retrieved.csv), II) count of retrieved species (species_retrieved.csv), III) count of repositories from which the records stem from (repositories_employed.csv), IV) count of retrieved genes per species (species_genes.csv), V) parsed information of each record (records_info.csv), and VI) summarized information about all the metadata metrics, including warnings if records have missing information. In those tables, since each record can be associated with multiple species and/or repositories, the count of these features may exceed the total count of records. An extra text file (VII) is also generated if a gene list is input to build the database, with all targeted genes that were not found after the download step (genes_not_retrieved.txt). Additionally, some metadata information is plotted to visualise the presence (species_per_gene_grid.png) and abundance (species_per_gene_barplot.png) of each gene per species **(Figure 2)**. Databases with more than 15 species or 35 genes (arbitrary criteria) might overload the plot information, making them unintelligible. In this case, a warning message is printed in the terminal.

**Figure 2.**
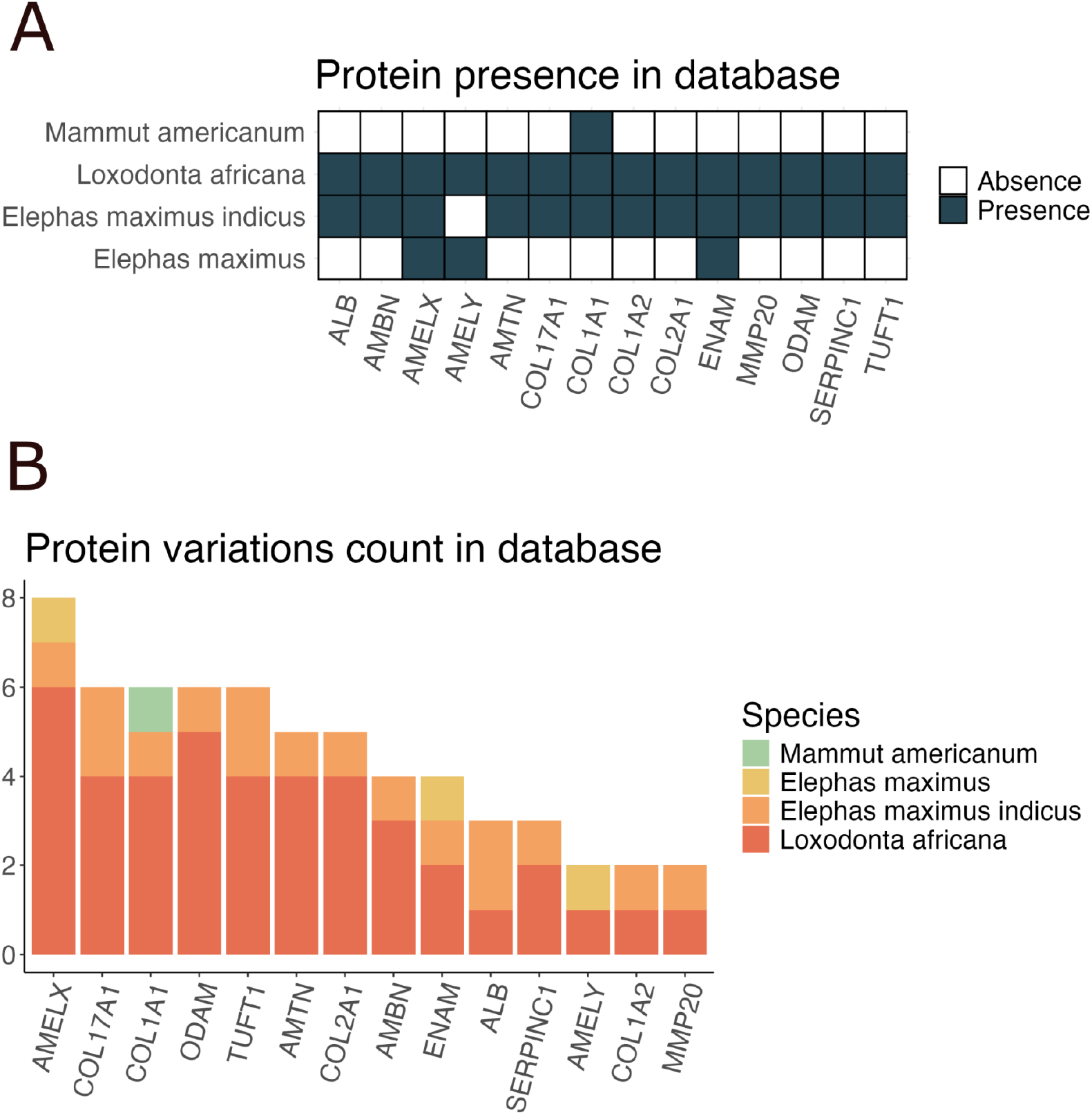
Metadata output plots to visualise the protein content and counts of a Proboscidea enamelome database (TaxID: 9779; gene list: documentation/example/enamelome.txt. **A)** Grid plot showing the presence or absence of a gene for each species. **B)** Barplot showing the number of different protein records (i.e. isoforms or protein variants) per species and gene. If a record is associated with multiple species, it is counted more than once. As a consequence, counts between species might overlap some records, eliminating the one-to-one correspondence between records and species per gene counts.

A step-by-step tutorial can be found in ProteoParc’s ‘documentation’ folder on GitHub. Additionally, two demo output results are stored there, consisting of a whole proteome download of the *Mammuthus* genus and an enamel tissue-specific (enamelome) download of the Proboscidea order. The enamelome gene list (example/enamelome.txt), employed to target the Proboscidea download, contains 15 usually detected enamel and dentine proteins reported in **Taurozzi et al., 2024**.

### Execution and Time Performance

Five scenarios were selected to test the performance of ProteoParc, covering different orders of magnitude in terms of records downloaded: Proboscidea enamel proteome (59 records ∼ 10^1^, TaxID: 9779); human enamel proteome (240 records ∼ 10^2^, TaxID: 9606); *Mammuthus* whole proteome (1249 records ∼ 10^3^, TaxID: 37348); mammalian enamel proteome (15544 records ∼ 10^4^, TaxID: 40674); and Proboscidea whole proteome (133166 records ∼ 10^5^, TaxID: 9779). For each scenario, all processes – *Download, Remove redundancy, Align* and *Metadata*– were consecutively run on a cluster node 2 x AMD EPYC 7H12 64-Core processor with a mean download speed of 50 MB/s. Each test was run 20 times to compute the mean execution *real* time through the bash *time* command. All the “enamel proteome” tests were generated by inputting an enamelome gene list.

ProteoParc’s execution time significantly increases with the number of records downloaded. This time increment is mostly due to the *Remove redundancy* and *Align* processes **(Figure 3)**, which recursively compare all the sequences present in the database. Specifically, the *Align* process is the main driver of computational time, while the *Remove redundancy* process influences the entire execution time only when downloading more than 10^5^ records. The execution time for gene-targeted downloads (scenarios 10^1^, 10^2^, and 10^4^) and small proteomes (scenario 10^3^) took less than 5 minutes in each process. Those results reflect realistic scenarios as the pipeline is intended to build tissue-specific (narrow) protein databases. Even though the 10^5^ scenario run took more than four hours to finish, the execution time is still feasible for an HPC job execution. Thus, the *Align* step might be omitted to remarkably reduce computational time in download scenarios with more than 10^5^ records, also considering that manually checking that many records is unfeasible. The *Remove redundancy* step might be deactivated as well, but then repetitive sequences will be present in the database, possibly compromising downstream analysis.

**Figure 3.**
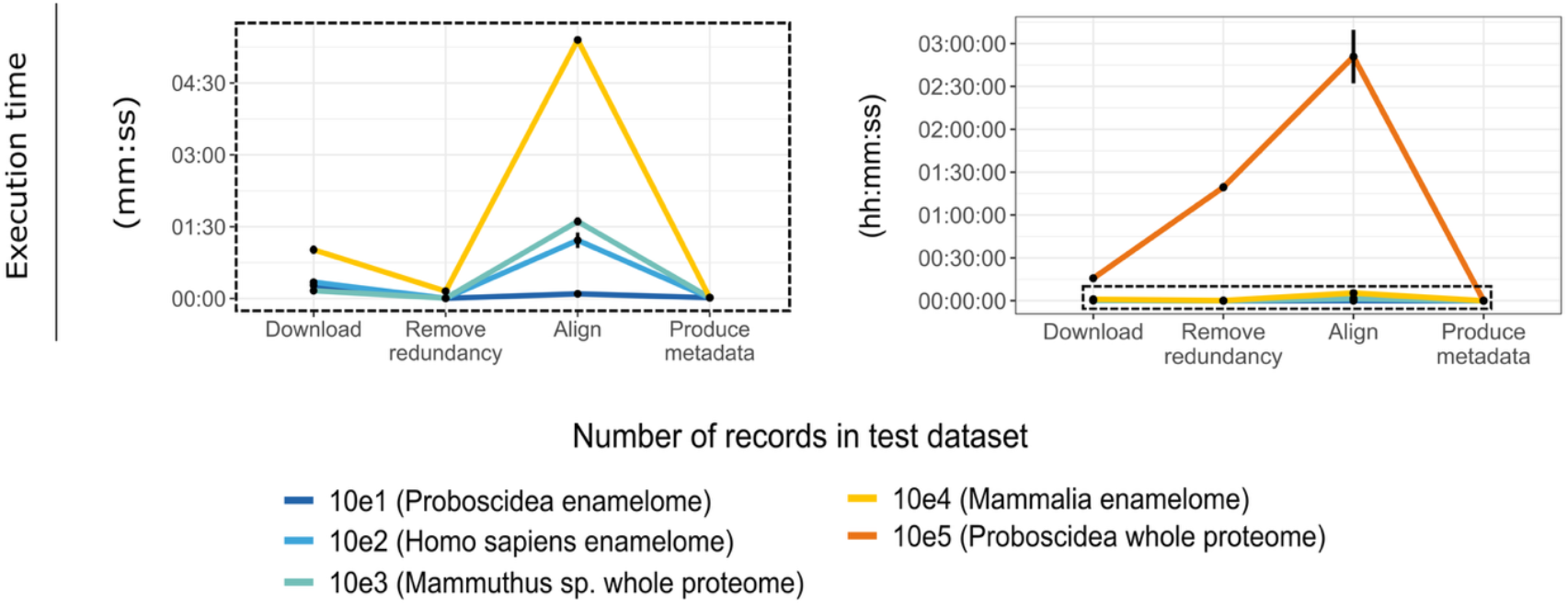
ProteoParc’s execution time in five different download scenarios: Proboscidea enamel proteome (59 records ∼ 10^1^, TaxID: 9779); human enamel proteome (240 records ∼ 10^2^, TaxID: 9606); *Mammuthus* whole proteome (1249 records ∼ 10^3^, TaxID: 37348); mammalian enamel proteome (15544 records ∼ 10^4^, TaxID: 40674); and Proboscidea whole proteome (133166 records ∼ 10^5^, TaxID: 9779). All processes were consecutively run 20 times to calculate the mean execution time on a node 2 x AMD EPYC 7H12 64-Core processor with a mean download speed of 50 MB/s. The standard deviation range is plotted at all points, but it only shows when the values are high enough to appear in the plot scale. The left plot is a zoom in from the right plot (dashed area), removing the 10^5^ scenario. The mean execution time grows exponentially with the number of proteins downloaded, and the *Align* process is the most time-consuming under all scenarios. In the 10^5^ scenario, the *Remove redundancy* process also plays a significant role in the whole computational time.

## Conclusion

Current bioinformatic tools employed in ancient and non-model organism studies have been designed to perform high-quality sample proteomics, but modified in a certain way to fit low-quality scenarios. Therefore, specific software still needs to be developed to assist in quality control, validation, and analysis of ancient and non-model proteomics. Software and pipelines such as deamiDATE **(Ramsøe et al., 2020)**, PaleoProPhyler **(Patramanis et al., 2023)** or Anubis **(Chiang, Nair, et al., 2024)** are good examples of helpful bioinformatic tools in these contexts. ProteoParc follows this trend, providing the community with a user-friendly and versatile tool to build reference databases for palaeoproteomic and non-model organism studies. It also removes format incompatibilities and provides metadata, allowing for database testing within different analyses. The option to build non-redundant and gene-specific databases will reduce computational costs and yield statistically more confident peptide sequences in MS/MS identification software. Moreover, custom databases can also behave as a starting point to be extended using newly translated sequences from genomic data. Thus, ProteoParc aims to help in one of the first critical steps of ancient protein data analysis, making research more reproducible and comparable in the newly developed field of paleoproteomics and non-model organisms analyses.

## Data Availability

ProteoParc: github.com/guillecarrillo/proteoparc

Execution time data: github.com/guillecarrillo/execution_time_proteoparc

## Acknowledgements

**G.C.M**. is supported by the programa predoctoral AGAUR-FI ajuts (2024 FI-1 00211) Joan Oró from Secretaria d’Universitats i Recerca, Departament de Recerca i Universitats de la Generalitat de Catalunya and the European Social Found Plus. **J.K**. is supported by the European Union’s Horizon 2020 research and innovation program under the Marie Sklodowska-Curie “PUSHH” training network, grant agreement No. 861389. **T.M.B**. is supported by funding from the European Research Council (ERC) under the European Union’s Horizon 2020 research and innovation programme, grant agreement No. 864203 and PID2021-126004NB-100 (MICIIN/FEDER, UE). This work is part of R+D+I projects PID2020-116908GB-I00 and PID2020-117289GB-I00, funded by the Agencia Estatal de Investigación of the Spanish Ministerio de Ciencia e Innovación (MCIN/AEI/10.13039/501100011033/). Research has also been supported by the Agència de Gestió d’Ajuts Universitaris i de Recerca of the Generalitat de Catalunya (2001 SGR 00620). We thank Ricardo Fong-Zazueta, Amanda Gutierrez, Joseph D. Orkin, Joanna L. Kelley, and Luis Ferrández for their feedback throughout the development stage.

## Author Contributions

Conceptualisation: E.L. and G.C. with assistance and guidance from all co-authors. Programming and analyses: G.C. Writing: G.C. with input from all co-authors.

